# Single-Molecule Methods to Investigate Mechanisms of Transcription by RNA Polymerase of *Mycobacterium tuberculosis*

**DOI:** 10.64898/2026.03.27.714832

**Authors:** Wenxia Lin, Omar Herrera-Asmat, Alexander B. Tong, Tiantian Kong, Carlos Bustamante

## Abstract

Single molecule methods have become prevalent tools in elucidating molecular processes across various life science fields over the past three decades, driving breakthroughs in understanding their underlying molecular mechanisms. In our study, we employed two single-molecule methods, Förster Resonance Energy Transfer (smFRET) and high-resolution optical tweezers, to investigate the transcription of *Mycobacterium tuberculosis* RNA polymerase (MtbRNAP) from initiation through to termination. We aim to provide a set of comprehensive biophysical tools to deepen our current understanding of MtbRNAP and its transcription factors. These experimental assays represent an important step towards unraveling the molecular dynamics and interactions that support transcription in *Mycobacterium tuberculosis*.

## 1. Introduction

*Mycobacterium tuberculosis* (Mtb) is the causative pathogen of tuberculosis (TB), the leading infectious disease killer worldwide, with 1.23 million deaths in 2024^1^. *Mycobacterium tuberculosis* RNA polymerase (MtbRNAP) is a conserved and essential enzyme that plays a crucial role in the adaptation of the pathogen during pathogenesis^2–4^ and, therefore, is a common target for anti-TB drugs. The transcription regulation of Mtb involves novel transcription factors and unique DNA elements that enable survival under host-induced stress; these factors are not present in the regulation of the more commonly studied *E*.*coli* RNA polymerase (EcoRNAP) system. Investigating the molecular mechanism of transcription in Mtb enhances fundamental biological understanding of this process and provides insights into developing strategies to inhibit it.

Earlier studies of MtbRNAP have primarily been structural, including the discovery of transcription initiation transition states as well as anti-TB drug and transcription factors binding sites ^5–14^. However, studies of the dynamics of MtbRNAP transcription remain largely unexplored, and recent single-molecule studies of MtbRNAP have begun to shed light on revealing the dynamics of this enzyme^15,16^.

Single-molecule methods can capture the dynamics of transient intermediates and rare states adopted by RNA polymerases during elongation, an observation of great interest as recent studies suggest that MtbRNAP pausing during transcription elongation enables Mtb to adapt to environmental changes, making the pausing phase a promising target for TB therapeutics^17^.

Single-molecule methods such as Förster Resonance Energy Transfer (smFRET), optical tweezers, and magnetic tweezers have been successfully used in the characterization of the molecular dynamics of other RNA polymerases, including EcoRNAP and yeast RNA polymerase II^18–24^. These methods showed promise for application to MtbRNAP as well. One previous smFRET study of MtbRNAP revealed single molecular dynamics of transcription activation in Mtb^15^. Our earlier research using high-resolution optical tweezers to study single-molecule dynamics of MtbRNAP demonstrated how various drugs differently affect transcription elongation^16^. In this paper, we employed a combination of single-molecule methods to study transcription initiation, elongation, and termination of MtbRNAP *in vitro*. Accordingly, we identified key biological questions, proposed experimental designs to address them, and presented preliminary data findings, shedding new light on understanding the molecular mechanisms of transcription in Mtb.

## 2. smFRET investigation of dynamics between MtbRNAP and transcription factors MtbCarD and MtbGreA during transcription initiation

In Mtb, transcription factors such as MtbCarD, RbpA, and sigma factors assist MtbRNAP to melt the DNA promoter, stabilize the open promoter complex, and slow down promoter escape^5,15,25–27^. These findings prompt the question of how MtbRNAP effectively proceeds from transcription initiation to transcription elongation, as well as what is its mechanism of regulation. MtbGreA, another conserved and essential mycobacterial factor, rescues backtracked and paused MtbRNAPs by promoting the endonucleolytic cleavage of nascent RNA during transcription elongation, a function common to Eco Gre factors^28–31^. However, MtbGreA has also been found to halve the amount of abortive transcripts and enhance promoter clearance during transcription initiation in Mtb^32^. While both MtbCarD and MtbGreA interact with the transcription initiation complex in Mtb, MtbGreA remains bound to the enzyme throughout transcription elongation, whereas MtbCarD dissociates during promoter clearance^33^. These observations raise several questions: Is MtbCarD dissociation from the transcription initiation complex influenced by MtbGreA? Do MtbCarD and MtbGreA jointly control the transition from transcription initiation to elongation? To address these questions, we aim to investigate potential interactions between these two transcription initiation factors.

Since smFRET is a powerful tool for exploring protein complex interactions within a spatial resolution of 3-8 nm^34^, it can be adapted to detect single-molecule dynamics and protein conformational changes between the MtbRNAP and multiple transcription initiation factors, including MtbCarD and MtbGreA. Therefore, we designed an smFRET assay in which the MtbRNAP transcription initiation complex, pre-assembled with a short DNA template in the presence of acceptor Cy5-labeled MtbCarD, is immobilized on a glass coverslip. Donor Cy3-labeled MtbGreA is then introduced to detect smFRET signals between the paired dyes and evaluate potential dynamic interactions between MtbCarD and MtbGreA (Fig. 1a). The specific protocols and preliminary results are described below.

**Fig. 1.**
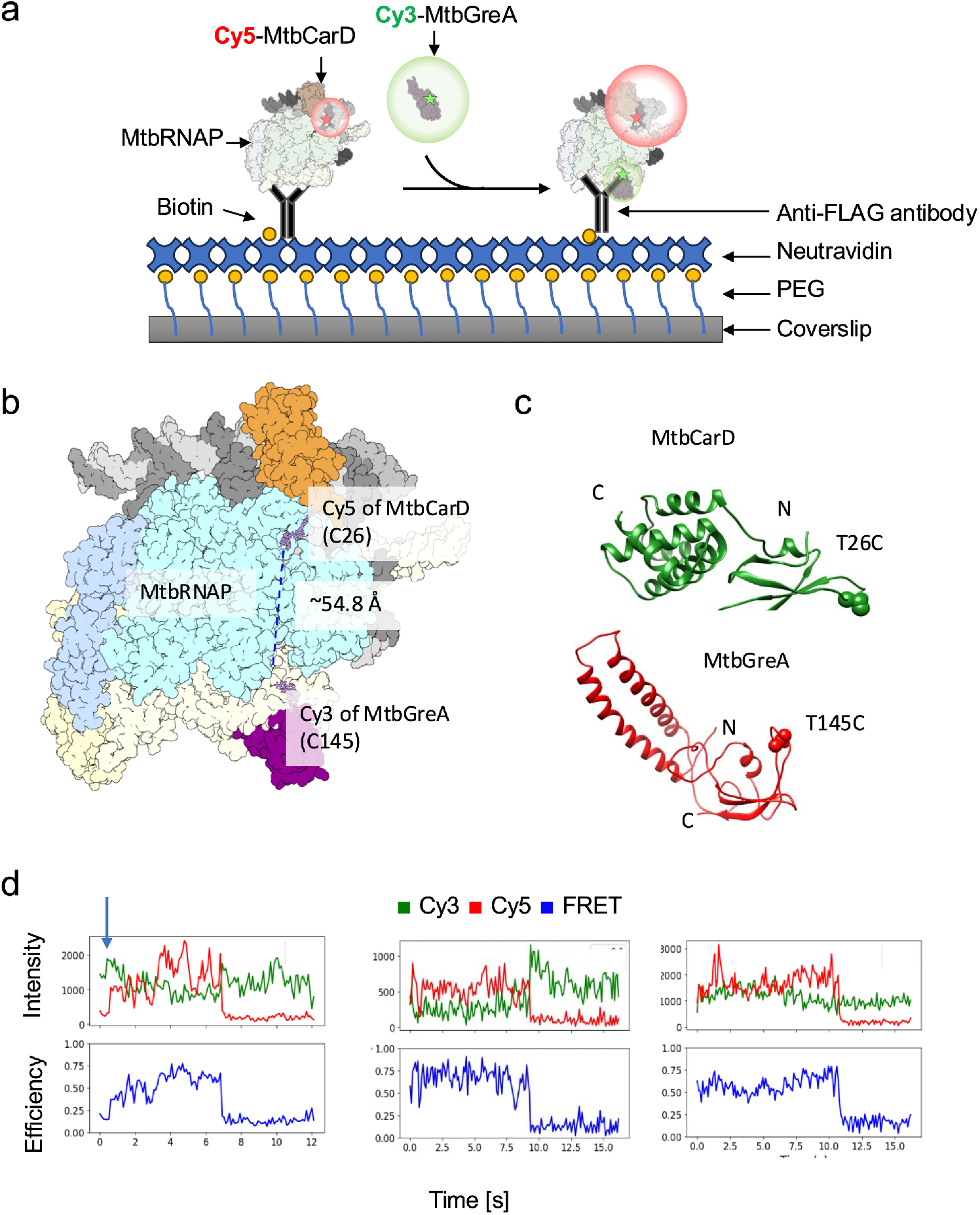
SmFRET investigation of dynamics between MtbRNAP and transcription factors MtbCarD and MtbGreA during transcription initiation. **a**. Illustration of smFRET assay design to study potential dynamic interactions between transcription factors MtbCarD and MtbGreA during transcription initiation. **b**. Fluorescent labeling sites of MtbCarD and MtbGreA are shown. The structural models of T26C MtbCarD and T145CMtbGreA were generated using PyMOL, based on chain J of PDB: 6EDT and a homology model derived from chain C of PDB: 6RIN, respectively. **c**. Predicted Cy3-Cy5 inter-dye distance withing the MtbRNAP complex, calculated from the structural alignment of the chains of the PBDs described in b, with Cy3 and Cy5 maleimide groups modeled onto the respective cysteine residues using custom SDF files. **d**. smFRET fluorescence data showing FRET between Cy5-MtbCarD and Cy3-MtbGreA.

### 2.1 Fluorescence labeling of MtbCarD and MtbGreA

The labeling strategy for MtbCarD and MtbGreA must maintain a donor-acceptor distance of 3-8 nm while not affecting their activity. A structural alignment was performed in Chimera^35^, aligning PDB 6EDT (MtbRNAP open promoter complex with MtbCarD and AP3 promoter)^5^ with a homology model of MtbGreA, built using PDB 6RIN (EcoRNAP backtracked elongation complex bound to GreB)^36^ via SWISS-model^37^(Fig. 1b). Using this model, we selected the two labeling sites as MtbCarD T26 and MtbGreA T145, as they are the right distance apart, are likely to tolerate cysteine substitutions for labeling, and are located in solvent-accessible areas and would likely accommodate the dyes (Fig. 1b, c). The calculated dye separation distance of 54.8 Å should yield a measurable FRET signal with the Cy3-Cy5 FRET pair of 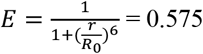.

MtbCarD T26C and MtbGreA T145C were purified using nickel columns with saline buffer (50 mM Tris-HCl, pH 8.0, 250 mM NaCl) with 1 mM DTT to prevent dimer formation. Histidine tags were removed through TEV protease cleavage, followed by reverse nickel column purification. Size exclusion chromatography (SEC) was then performed using the saline buffer containing 5 mM DTT to isolate the monomeric forms of the proteins. The proteins were further purified with ammonium precipitation. Labeling was done using maleimide chemistry^38,39^. Free dyes were removed by capturing the proteins with HiTrap Q HP (Cytiva) in buffer TGED-200 (20 mM Tris-HCl, pH 7.9, 0.1 mM EDTA, 1 mM EDTA, 5% Glycerol, and 200 mM NaCl) and eluting them in a linear gradient with buffer TGED-1000 (100 mM NaCl). The proteins were concentrated, and then the buffer was exchanged to TGED-400 (400 mM NaCl) via centrifugation. The labeling yields were above 75% as determined by UV-Vis spectrophotometry.

The activity of Cy5-MtbCarD was verified by showing that it was able to form a transcription initiation complex with MtbRNAP in an electrophoretic mobility shift assay (EMSA) (Fig. S1a). For Cy3-MtbGreA, we used a bulk nucleotide misincorporation assay and showed that it reduced misincorporation at a rate comparable to the wild-type enzyme (Fig. S1b).

### 2.2 Transcription initiation open complex preparation

For initiation studies, a FLAG tag was introduced at the C-terminus of the β’ subunit of the polymerase to enable surface immobilization through a biotinylated mouse anti-FLAG (FG4R) antibody. A TEV protease site was also placed after the His tag of the alpha subunits to remove any tag that can affect transcription initiation. The purification of the polymerase as a holoenzyme with σ^A^ was conducted according to established protocols^40^, with an additional step of TEV proteolysis and reverse nickel chromatography between the nickel and MonoQ chromatography steps to remove the His tag.

The 290 bp DNA template, which contains the promoter AC50, a C-less cassette, and a fragment of the Mtb gene rpoB, was obtained by PCR from plasmid pUC-del-LAC-AC50-MtbrpoBC, which was previously used to obtain templates for single-molecule assays of MtbRNAP transcription elongation (SI Table 1)^16^.

Transcription initiation open complexes were assembled by incubation of 20 nM FLAG-tagged holoenzyme-σ^A^, 200 nM Cy5-MtbCarD, and 200 nM 290 bp DNA template containing AC50 promoter at 22 °C for 10 mins, followed by addition of 70 nM of biotinylated anti-FLAG antibody and additional incubation of 10 min before storage on ice. This transcription initiation open complex was named bio-Ab-TIC-Cy5-CarD. The bio-Ab-TIC-Cy5-CarD complexes were used within one hour to ensure complex integrity.

### 2.3 smFRET assays and data acquisition

smFRET assays were performed in microfluidic chambers made by commercial biotin/PEG glass coverslips and cleaned quartz slides subjected to quality control of the number of fluorescent dust spots (Fig. 1a). Microfluidic chambers were prepared by first allowing for equilibration in buffer, passivation with 30 mg/mL beta-casein, coating with 3 µM neutravidin, and finally introduction of 2.7 nM bio-Ab-TIC-Cy5-CarD. Each step was done for 10 minutes at 22 °C in TB130 buffer (20 mM Tris-HCl, pH 7.9, 130 mM KCl, 10 mM MgCl_2_, 5 mM NaN_3_ and 1 mM DTT), with three washes of plain TB130 between incubations. Experiments were performed on a home-built objective-based TIRF microscope at 22 °C. The immobilization density of bio-Ab-TIC-Cy5-CarD complexes was verified through Cy5 excitation by a red laser under an imaging buffer (TB130 supplemented with 2 mM Trolox, 50 mM Glucose, 12.5 mM L-Ascorbic Acid, 1 mg/ml Casein, 0.9 U/ml Pyranose oxidase, 0.25 mg/ml Catalase, and 0.2 U/µl RNaseOUT). Afterwards, Cy3-MtbGreA was flowed into the chamber with imaging buffer, and recordings were taken with exposure times ranging from 33.3 ms to 100 ms, using excitation from a diode-pumped 532-nm laser. Fluorescence emission was detected using an EMCCD camera (IXon EM+ 897; Andor).

The magnitude of Cy3-MtbGreA concentration was diluted to a range of 4-10 nM to have acceptable fluorescence background.

### 2.4 smFRET time trajectories and data analysis

FRET efficiency was calculated with the equation E = I_A_/(I_D_ + I_A_), where I_A_and I_D_ respectively correspond to acceptor and donor fluorescence intensities after correction for background and leakage from Cy3 into the Cy5 channel. Recorded smFRET time trajectories revealed the dynamical interactions between Cy5-MtbCarD and Cy3-MtbGreA (Fig. 1d). Some trajectories also captured the initial binding steps of Cy3-MtbGreA or its interactions with bio-Ab-TIC-Cy5-CarD (Fig. 1d, see the arrow).

When the Cy5 fluorescence intensity decreased to zero, several possibilities were considered: (a) bleaching of the Cy5 dye, (b) MtbCarD staying bound but moving further away from MtbGreA, or (c) MtbCarD dissociating from the transcription initiation open complex. However, we currently lack evidence to determine which of these possibilities occurred.

## 3. Optical tweezers analysis of the effect of MtbCarD and MtbGreA on transcription elongation by MtbRNAP

Mtb was found to only have a single GreA factor, in contrast to Eco, which harbors two Gre factors, GreA and GreB, along with additional secondary channel binding proteins to compensate for Gre function^41,42^. EcoGreA cleaves 2-3 nucleotides, whereas EcoGreB cleaves 9 nucleotides^43^. MtbGreA generates cleavage products primarily consisting of 2-3 nucleotides, similar to the activity of EcoGreA^32^. MtbGreA performs a crucial role in rescuing paused and arrested MtbRNAP on GC-rich templates during transcription elongation, and it has been revealed to be essential for cell growth and survival, facilitating the organism’s adaptation to adverse stresses that threaten its viability^32^. Knocking down Gre factor in *Mycobacterium smegmatis* led to altered expression of various topology and transcription regulators^44. 32^

We utilized an optical tweezers assay to study transcription elongation by MtbRNAP, comparing its behavior with and without MtbGreA. Additionally, we conducted transcription elongation experiments in the absence and presence of MtbCarD, a highly expressed essential factor and global regulator for mycobacterial pathogenesis^45–47^. As noted earlier, MtbCarD has not been observed during the transcription elongation phase and is not expected to interfere with MtbGreA function. The optical tweezers geometry is shown in Fig. 2a, in which the biotinylated MtbRNAP is halted on the C-less DNA template and attached to the DNA handle via a biotin and neutravidin link. Both the DNA handle and the DNA template were linked to microbeads, which were held in two optical traps. As transcription occurs, the length of the DNA tether decreases, and the tweezers system moves the traps to keep the tension in the tether constant. This change in trap position over time is a measurement of transcription activity, reflecting the kinetics of nucleotide addition and polymerase progression^44^.

**Fig. 2.**
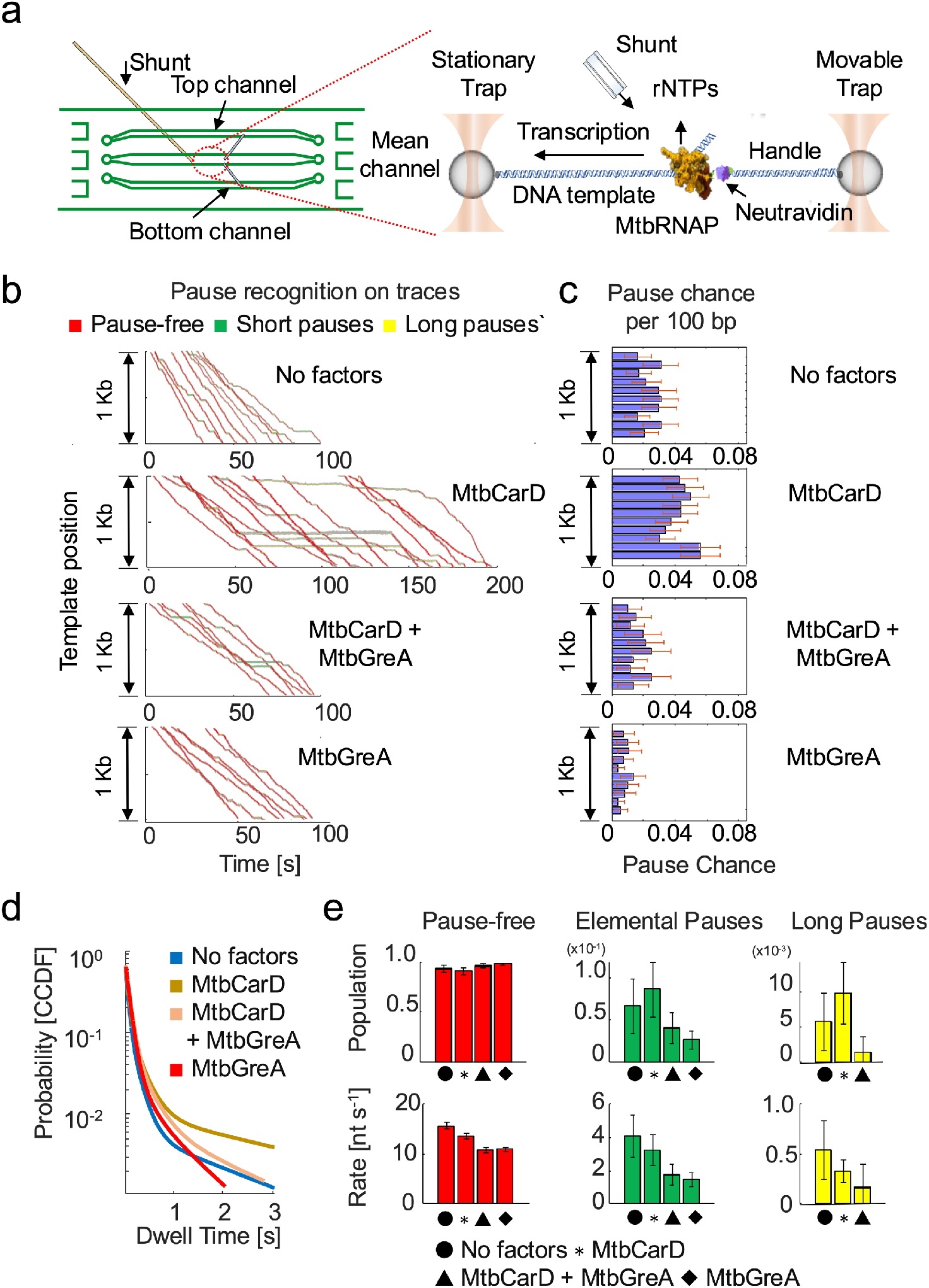
Optical tweezers investigation of the effect of transcription factors MtbCarD and MtbGreA on transcription elongation by MtbRNAP. **a**. Optical tweezers experiment to study MtbRNAP transcription elongation with/without transcription factor regulation. Left: schematic of the microfluidic chamber, illustrating the main, top and bottom channels as well as the shunt. Right: experimental geometry depicting the opposing force single molecule tether setup, in which a stalled MtbRNAP is used for transcription elongation measurements. **b**. Full single molecule trajectories of MtbRNAP transcription elongation under different conditions. **c**. Pause chance over 100 bp regions during transcription elongation under different conditions. **d**. Comparison of the complementary cumulative distribution function (CCDF) of the dwell time distributions (DTD) of MtbRNAP under the different elongation conditions. **e**. Population and rates obtained via DTD analysis from (d).

Below are the details about the transcription elongation experimental design, data analysis, and preliminary results.

### 3.1 DNA template and optical tweezer handle preparation

The digoxigenin-labeled 2.8 kb DNA template was made by PCR from plasmid pUC-del-LAC-AC50-MtbrpoBC, as shown in earlier research^16^.The DNA template contains the AC50 promoter, a transcription start site (TSS), a C-less cassette, and fragments of rpoB and rpoC genes in Mtb H37Rv, including the intergenic region. See 2.8 kb DNA template sequence in supplementary information (SI, Table 1).

Using PCR, we made a 1.5 kb DNA handle from a region of the Lambda genome without a promoter, labeling it with biotin on one end and double-digoxygenin on the other end.

### 3.2 Preparation of ternary elongation complexes

We started by combining 1 µM of biotinylated MtbRNAP with 10 µM of MtbCarD in a reaction volume of 2 µL at 37° C for 5 minutes. The role of MtbCarD in this procedure is to facilitate MtbRNAP to form the transcription initiation open complex. Afterward, we diluted the biotinylated MtbRNAP in TB40 buffer (20 mM Tris-HCl, pH 7.9, 40 mM KCl, 5 mM MgCl_2_, and 0.5 mM TCEP) to a final concentration of 10 nM. Next, we incubated 5 nM of the above transcription initiation open complexes with 150 µM of GpG and 2 nM of a digoxigenin-labeled 2.8 kb DNA template in a stall buffer (TB40 plus 3 µM of rATP, rGTP, and rCTP) and left it at 37°C for 20 minutes. Finally, we added 50 µg/ml of heparin and incubated at 22° C for 10-15 minutes to eliminate incompetent complexes. The ternary elongation complexes (TECs) were kept on ice before subsequent experiments. Note that the concentration of leftover MtbCarD from the transcription initiation open complex will be less than 13 pM in the optical tweezers assay.

### 3.3 Bead deposition

Anti-dig beads were passivated by vortexing with 2 mg/mL BSA for 5 min. Then, 0.1 nM of the TECs prepared above were mixed with 0.05 to 0.10% passivated anti-dig beads in buffer TB40 at 22°C for 10 minutes to deposit the complex on the beads. Separately, we combined 0.2-0.5 nM of biotin- and digoxigenin-labeled DNA handles with 300 nM Neutravidin and 0.05 to 0.10% passivated anti-dig beads in buffer TB40 at 22°C for 10 minutes to deposit the handles on the other beads. These coated beads can be preserved by flash freezing them for storage at −80°C or keeping them on ice for immediate use.

### 3.4 Optical tweezers assay to track transcription elongation in real time

µL each of DNA handle-linked beads and TEC-linked beads were diluted into 1 ml of degassed TB130 buffer and introduced into the top and bottom channels of a three-channel optical tweezers microfluidic chamber, in which dispenser tubes connect the top and bottom channels to the middle channel (Fig. 2a). Using two optical traps, we captured one of each bead in each trap, relocated them to a clear, undisturbed area, and brought the beads in proximity to each other to form the linkage between the biotin on the TEC and the neutravidin on the DNA handle. The beads were moved away from each other to check for the formation of a tether, and the observed length of the tether was used to determine whether a single tether or multiple tethers were formed. Immediately after forming a single-molecule tether, we set a constant force of 4-5 pN, recorded the distance between two beads, and opened a shunt to introduce 1 mM NTPs to resume transcription elongation, supplemented with 1.5 µM MtbCarD and/or 0.5 µM MtbGreA.

### 3.5 Single molecule full trajectories and quantitative data analysis

Fig. 2b presents single-molecule trajectories of MtbRNAP transcription elongation, highlighting pause-free periods (red) and interspersed pauses (yellow-green). We found that adding excess MtbCarD during transcription elongation increases the pause density, as indicated by more frequent pauses in comparable DNA regions across multiple traces (Fig. 2c). In contrast, MtbGreA reduced the MtbCarD-induced pauses (Fig. 2b, 2c). This result for MtbCarD is interesting given the *in vivo* findings that it is primarily associated with initiation^47,48^.

A quantitative analysis of transcription elongation trajectories was conducted to distinguish different states (on-pathway, short pause, long pause, etc.) of nucleotide addition^49^. The transcription traces were fit to a monotonic 1 bp staircase to extract dwell times. The CCDF of these dwell times is shown in Fig. 2d^49^. The dwell time at each nucleotide was fit using a sum of exponentials: 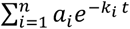, where *i* represents the dynamic state of MtbRNAP, *a*_*i*_ is the state probability (subject to the constraint ∑_*i*_ *a*_*i*_ = 1), and *K*_*i*_ is the kinetic rate associated with state i, ordered in descending value (k_1_ > k_2_ > … > k_n_). This model enables the analysis of distinct nucleotide addition states. Based on this analysis, MtbGreA was found to globally modulate MtbRNAP activity by reducing the duration and frequency of pauses and the elemental pause rate even in the presence of MtbCarD (Fig. 2e). The elemental pause rate was associated with the rate of escape by MtbRNAP from elemental pauses.

## 4. Optical tweezers investigation of RNA co-transcriptional folding during transcription termination

Regarding transcription termination by MtbRNAP, recent findings indicate that nearly 90% of terminator structures in Mtb lack a U-tract or contain a mixed U/A-tract^50–53^. This observation presents a noteworthy deviation from classical models of transcription termination, mostly based on studies of EcoRNAP. The classic transcription termination model relies on two elements: a terminator hairpin structure in the form of a stem-loop and an adjacent U-tract^54,55^. The U-tract plays a role of creating a region of reduced base-pairing strength, which is essential for facilitating the efficient dissociation of the transcription complex^56,57^. This model may not be a perfect fit for MtbRNAP. Another recent study has proposed that the poly U trail is not necessary for intrinsic transcription termination in Mtb and other bacteria^53^. Given that many studies have documented the modulation of RNA polymerase transcription by RNA co-transcriptional folding^58,59^, an approach centered on the RNA co-transcriptional folding landscape presents an opportunity to gather direct observations that can be correlated with transcription termination, enabling a quantitative elucidation of the molecular mechanisms governing transcription termination by MtbRNAP. Furthermore, different termination efficiencies were observed in bulk between MtbRNAP wild-type and mutants^60^, which implies potential differences in their co-transcriptional folding landscapes.

A protocol is presented for capturing real-time RNA co-transcriptional folding trajectories employing time-shared dual optical tweezers. In this configuration, a TEC comprising a biotinylated wild-type MtbRNAP is stalled on a DNA template. The upstream end of the DNA template is anchored to an oligo-coated bead, and the nascent RNA strand from the TEC is captured through hybridization with a single-stranded overhang of a DNA handle attached to an anti-digoxigenin-coated bead (Fig. 3a shows the geometry). A similar optical tweezers geometry was previously designed to study signal recognition particle RNA^61^.

**Fig. 3.**
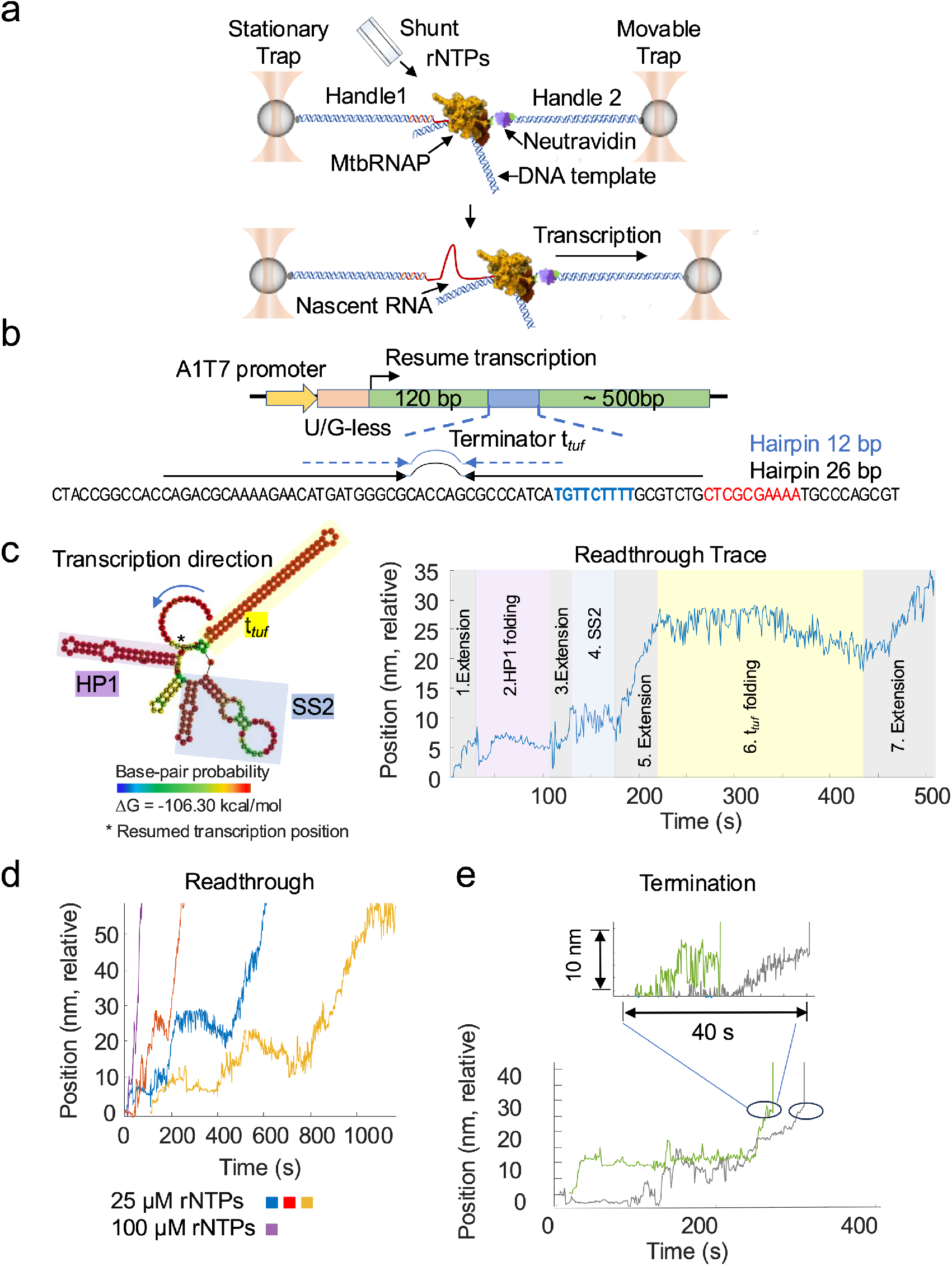
Optical tweezers investigation of RNA co-transcriptional folding during transcription termination. **a**. Optical tweezers geometry of RNA co-transcriptional folding by MtbRNAP during transcription termination. **b**. DNA template design for MtbRNAP termination. c. Left, RNAfold prediction of the folding of t_*tuf*_ RNA. The * show the position of resume of transcription elongation. Right, annotated readthrough trace showing the RNA extension and folding features. **d**. A collection of read-through trajectories obtained at varying [rNTP]. **d**. Termination trajectories.

### 4.1 DNA template design and preparation

The DNA template consisted of the A1T7 promoter, a U- and G-less cassette, a plasmid sequence spacer, the t_*tuf*_ terminator, and the remainder of the plasmid sequence. The t_*tuf*_ terminator was chosen to study MtbRNAP transcription termination due to its distinct structural features: a 12 bp RNA hairpin and an imperfect U-tract embedded within a DNA sequence predicted to form a longer hairpin of 26 bp (Fig. 3b). Experimental results indicate that the t_*tuf*_ terminator exhibits moderate termination efficiency (∼50%) *in vitro*^62^.

To produce the DNA template described above, we purchased a DNA encoding the promoter and the G&U-less cassette; we extracted the t_*tuf*_ terminator from Mtb via PCR and cloned them into the pJET1.2 plasmid (pMB1 family). This procedure resulted in the creation of the plasmid that contains the DNA template. The DNA template was obtained from a restriction digest of the pJET1.2 plasmid using NcoI and HindIII endonucleases. (See the whole DNA template sequence in SI, Table 1).

### 4.2 Biotinylated handle bead deposition

Approximately 250 pM of a 1.5 kb DNA handle (with biotin at one end and a 5’ P-ACCG overhang at the other) was incubated in TB40 buffer containing 0.1 mM ATP, 300 nM Neutravidin, 0.03% w/v 5’ P-CGGT oligo beads (1 µm, passivated with casein), and 40 U/µL of T4 DNA ligase at 22°C for 1 hour. Following this procedure, heparin was added to a final concentration of approximately 30 µg/ml, and the incubation continued for an additional 10 minutes.

### 4.3 TEC bead deposition

MtbRNAP TEC was made by mixing 20 nM of biotinylated polymerase with TB40 buffer that had 20 μM of rATP, 20 μM of rCTP, 150 μM of ApU, 20 nM of RNaseOUT, and 2 nM of DNA template. We maintained the incubation at 37°C for 15 minutes. Following this, heparin was added until it reached a final concentration of 50 µg/ml. It was then mixed with about 2 nM of a 3’ digoxigenin-labeled 1.5 kb DNA handle that had a 5’ overhang that is complementary (5’-TTGTGGTGGTTGTGGTGGTTGTGGTGGGAT) to the 5’ end of the nascent RNA transcript of the stalled TECs. We incubated this mixture for an additional 20 minutes at 22°C to eliminate nonfunctional complexes and capture the nascent RNA through in situ hybridization. We then added anti-digoxigenin (AD)-coated beads to the mixture and incubated it for an additional 20 minutes at 22°C.

### 4.4 Optical tweezers RNA-pulling assay for transcription termination

The two bead depositions were placed on ice and diluted to 1 ml with TB40 buffer containing 5 mM NaN_3_ separately. The suspensions were injected into two side channels of the microfluidic chamber, allowing the beads to diffuse into the central channel, where transcription resumed later. All experiments were conducted in this main channel. Using optical tweezers, we separately trapped the DNA handle bead and the stalled TEC bead. The beads were then brought together until a single tether formed. The coupled bead pair was moved to a clear area free of interference and a constant force of 6 to 9 pN was applied to the coupled bead pair. Transcription was restarted by adding rNTPs from 25 to 250 µM and the tether length was recorded over time.

### 4.5 Nascent RNA co-transcriptional folding trajectories observation and data analysis

The resulting trajectories have increases in extension consistent with transcription, interspersed with periods of rapid fluctuation attributable to RNA folding, as well as steady plateaus that indicate MtbRNAP pausing. In certain instances, abrupt terminations correspond to either transcriptional termination or tether breaking (Fig. 3c, d, and e). A key challenge in the analysis of co-transcriptional RNA folding trajectories is that, during ongoing transcription, the RNA tether becomes longer, whereas concurrent RNA folding results in a reduction of tether length.

To aid interpretation of the trajectories, we used RNA secondary structure prediction via the Vienna RNA package^63^. The folding of the RNA segment, spanning from the start of the G/U-less cassette to the loop of the t_*tuf*_ hairpin, was predicted, and the resulting secondary structure revealed three distinct features: a single hairpin (Hairpin 1, HP1), a set of three hairpins collectively identified as Secondary Structure 2 (SS2), and the t_*tuf*_ hairpin (t_*tuf*_) (Fig. 3c). The applied force of 6-9 pN is expected to not prevent the folding of these large structures but prevent the transient formation of smaller RNA secondary structures. Based on these predictions, the RNA co-transcriptional folding trajectories are anticipated to show the following events: extension → HP1 folding → extension → SS2 folding → extension → t_*tuf*_ folding, followed by either termination or readthrough. Extension changes corresponding to each RNA structural element are annotated on a representative readthrough trajectory in Fig. 3c. Based on these extension signatures, trajectories were categorized into two distinct classes: readthrough (Fig. 3d) and termination (Fig. 3e). These three readthrough trajectories each exhibited an initial rise, a stall at 5–10 nm, a noisy increase to 20–30 nm, gradual shortening to approximately 20 nm, and a subsequent noisy increase that concluded beyond 70 nm, corresponding to the transcription and folding of the predicted secondary structures (Fig 3c). Two termination trajectories were observed, which shared similar features at the start to the readthrough trace, except that the tethers are observed to break at 30nm extension in the t_*tuf*_ hairpin folding region. Tether breaking is the readout for transcription termination. In addition, several short trajectories were also observed (Fig. S2a), which likely resulted from premature tether loss due to mechanical rupture or RNAP dissociation before reaching the t_*tuf*_ region. Other trajectories may represent RNAP stalling or extended pausing; for instance, in one case, MtbRNAP transcribed slowly for over 1000 s following a ∼10 nm increase, although only ∼70 s was required to reach that point (Fig. S2b).

## 5. Concluding Remarks

These established methodologies provide a biophysical framework for investigating the molecular dynamics of transcription initiation, elongation, and termination in Mtb. smFRET trajectories involving Cy5-MtbCarD and Cy3-MtbGreA indicate a potential interaction between MtbCarD and MtbGreA during the transcription initiation phase. This observation may help elucidate the mechanism underlying MtbCarD dissociation at this stage, as our preliminary data do not provide definitive evidence that MtbGreA directly induces the release of MtbCarD in early transcription. Comprehensive analyses of single-molecule transcription trajectories elucidate the role of MtbGreA in transcription elongation by MtbRNAP. Notably, MtbCarD increases the frequency of pausing in MtbRNAP, whereas MtbGreA mitigates this effect. Overall, our results spanning both initiation and elongation phases indicate a potential global interference mechanism between MtbCarD and MtbGreA via allosteric binding to MtbRNAP.

The single-molecule termination assays provide quantitative measures of transcription rates and offer insights into RNA secondary structure formation before termination hairpin emergence. It is important to note that the RNA folding trajectory presented here is only a proposal, and additional truncation and mutagenesis experiments, guided by specific hypotheses, are needed to confirm features observed in RNA co-transcriptional folding trajectories and reinforce RNA folding predictions^59^.

In conclusion, although significant questions persist regarding transcription mechanisms in Mtb, single-molecule approaches yield valuable insights into MtbRNAP function and facilitate future exploration of its mutants, with implications for the development of novel drug strategies.

## Supplementary material

**Fig. S1.**
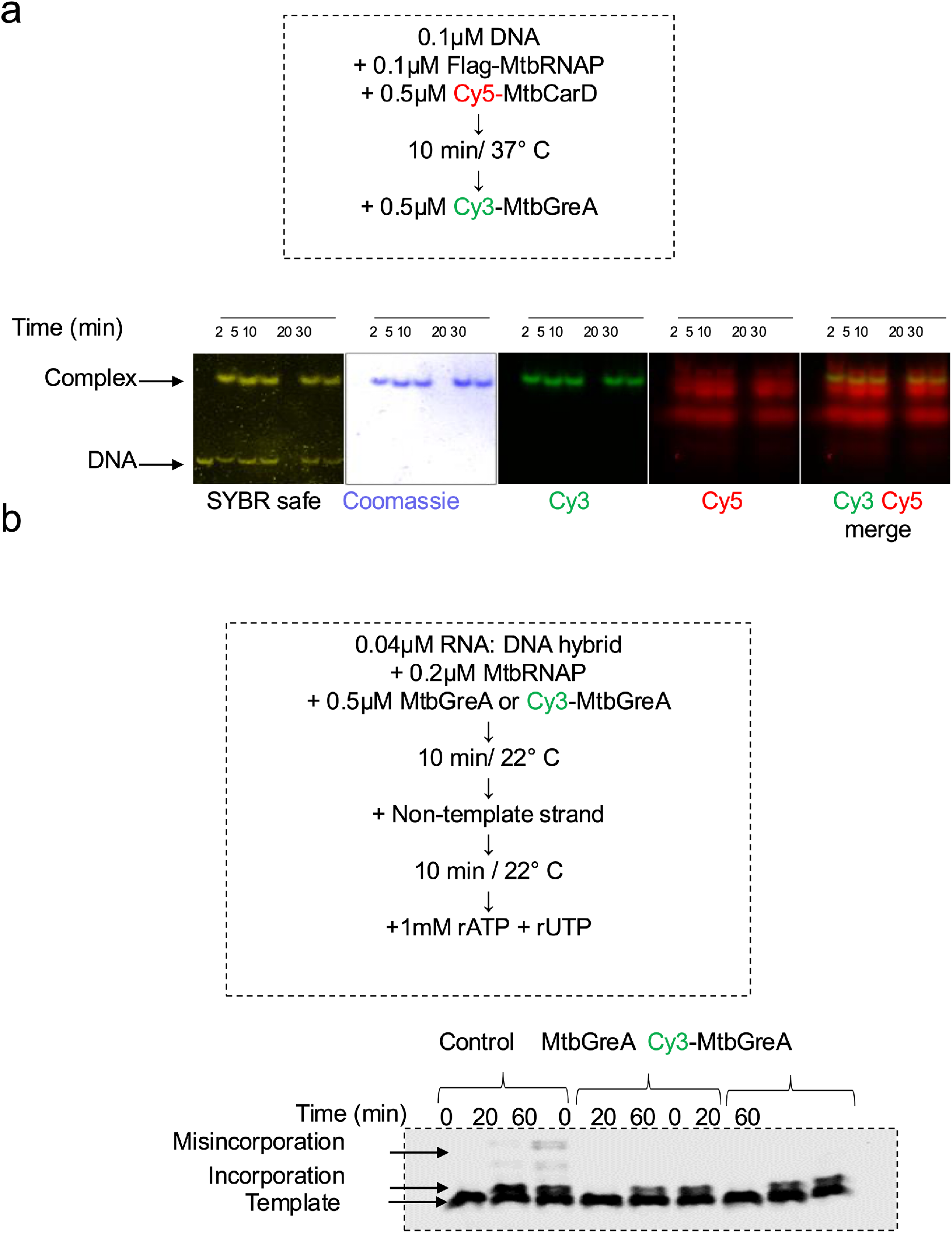
Evaluation of Cy5-MtbCarD and Cy3-MtbGreA functional activities. **a**. An electrophoretic mobility shift assay to verify a transcription initiation complex formation with Cy5-MtbCarD. **b**. A misincorporation bulk assay to verify Cy3-MtbGreA activity.

**Fig. S2.**
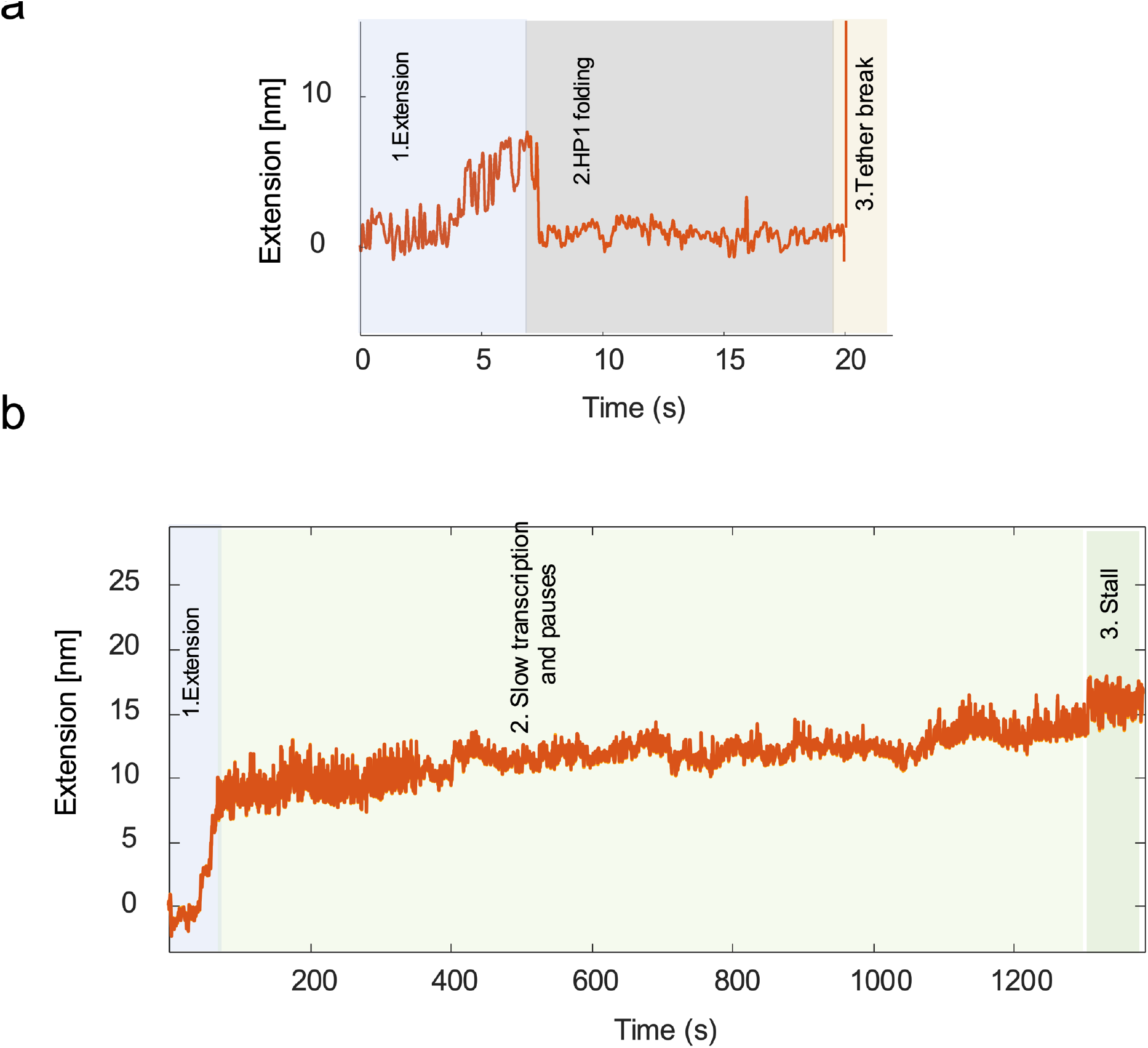
Other RNA co-folding trajectories. **a**. One shortened trajectory. **b**. One trajectory indicated that MtbRNAP paused, slowly transcribed, and finally stalled.

**SI. Table 1.**
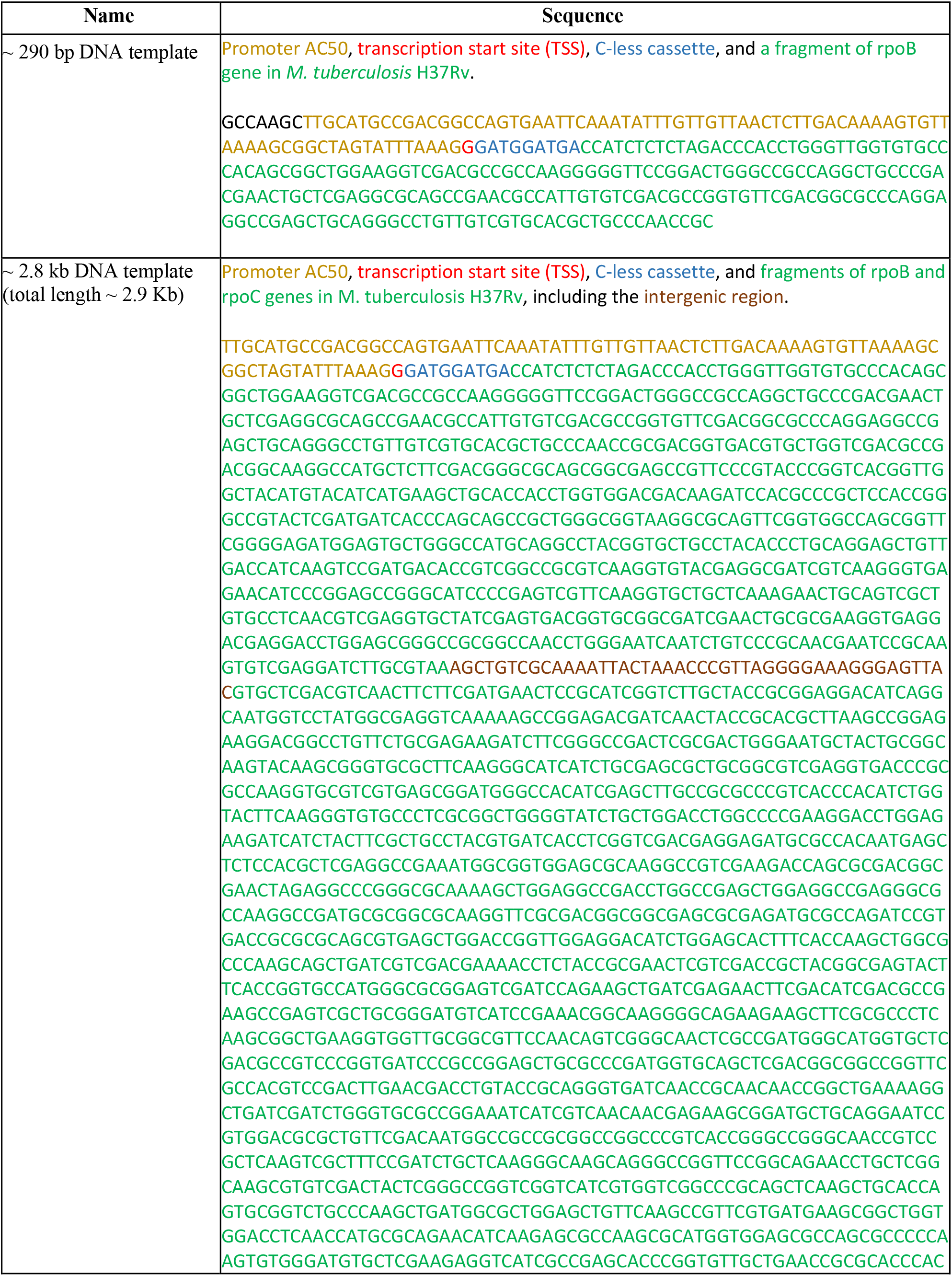
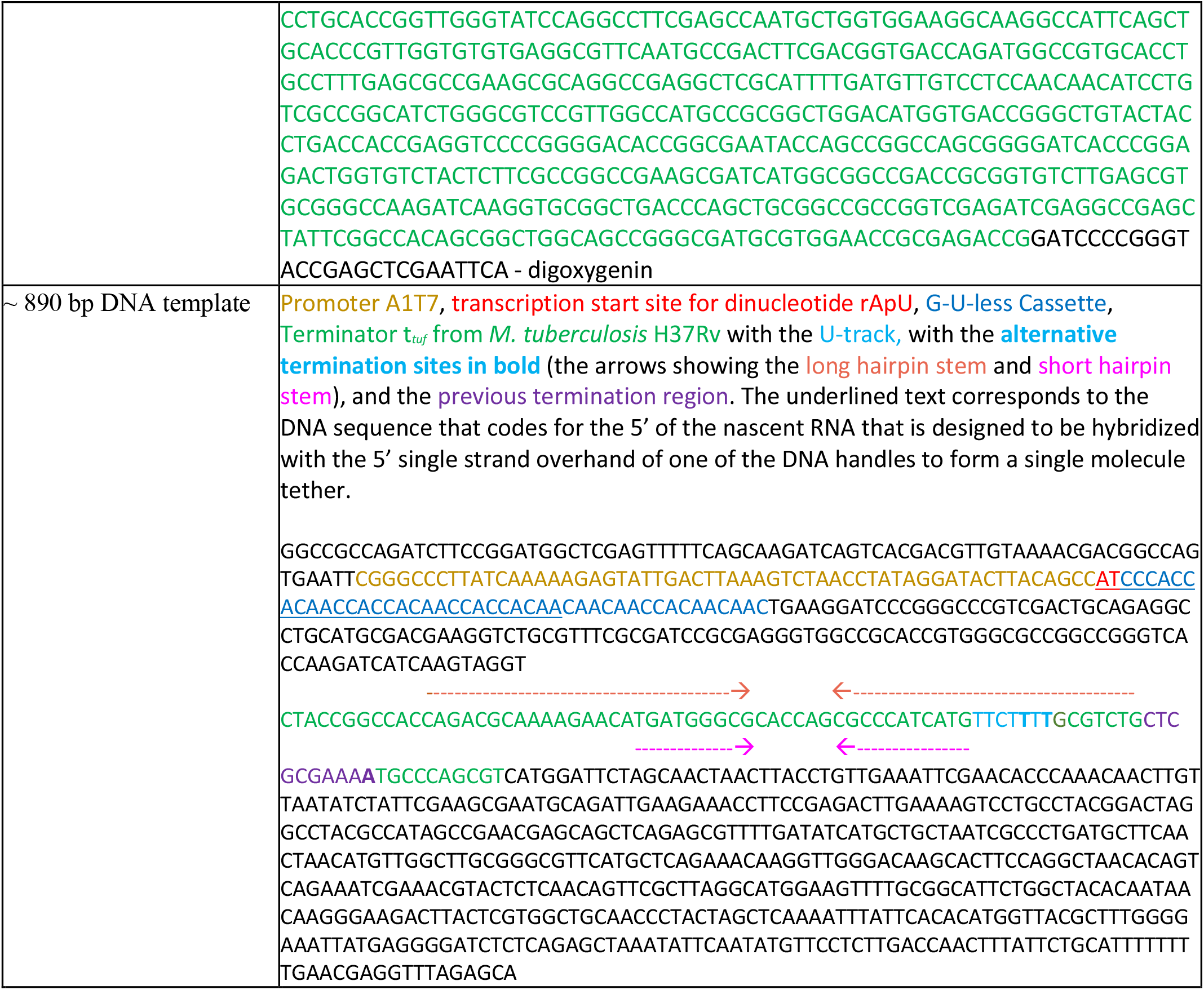

**SI. Table 2.**
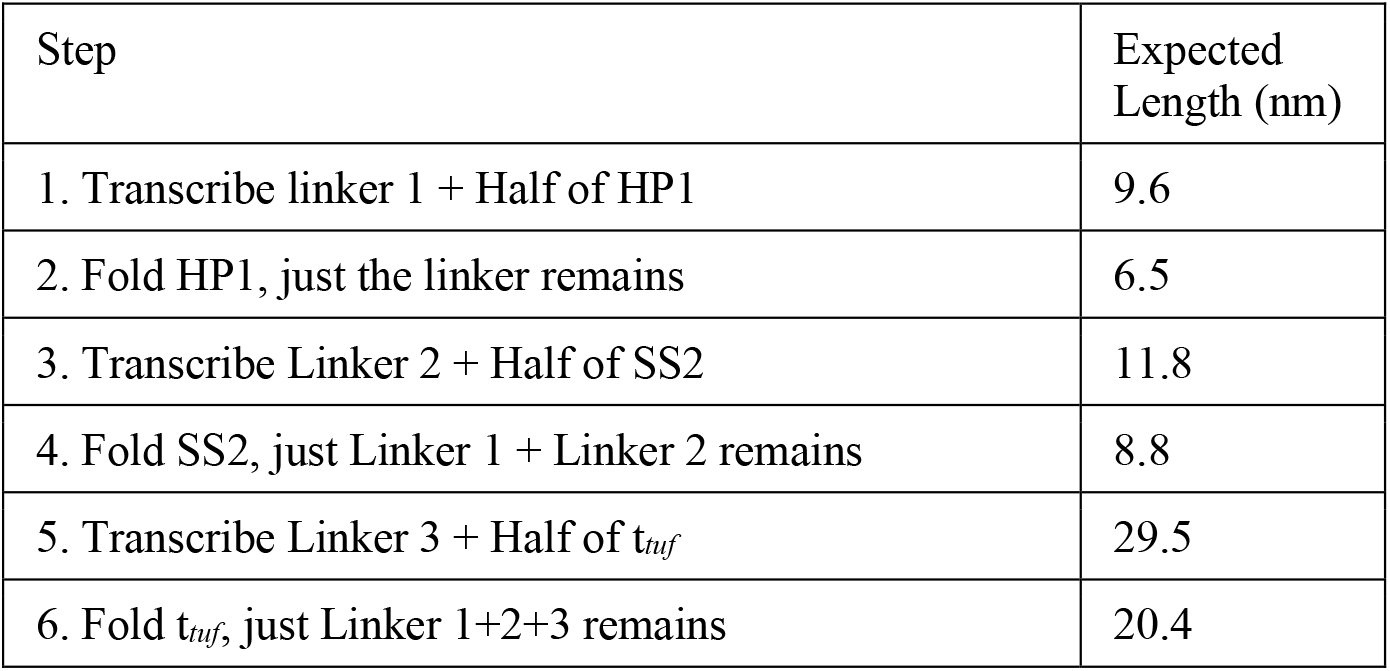

